# The effect of ascertainment on penetrance estimates for rare variants: implications for establishing pathogenicity and for genetic counselling

**DOI:** 10.1101/2023.02.17.528910

**Authors:** Andrew D. Paterson, Sang-Cheol Seok, Veronica J. Vieland

## Abstract

Next-generation sequencing has led to an explosion of genetic findings for many rare diseases. However, most of the variants identified are very rare and were identified in small pedigrees, which creates challenges in terms of penetrance estimation and translation into genetic counselling in the setting of cascade testing. We use simulations to show that for a rare (dominant) disorder where a variant is identified in a small number of small pedigrees, the penetrance estimate can both have large uncertainty and be drastically inflated, due to underlying ascertainment bias. We have developed PenEst, an app that allows users to investigate the phenomenon across ranges of parameter settings. We also illustrate robust ascertainment corrections via the LOD score, and recommend a LOD-based approach to assessing pathogenicity of rare variants in the presence of reduced penetrance.

---

Next-generation sequencing has led to an explosion in the number of genetic findings for many rare diseases. For certain types of rare coding variants (e.g. missense, or protein truncating), if the variant is sufficiently rare and has bioinformatic predictions that are severe, current algorithms result in it being classified as pathogenic (1). However, the analysis of large-scale sequencing from cohorts, such as ExAC (2), gnomAD (3), and the UK Biobank (4), has shown that many such variants may often lack clinically significant impact. For example, ExAC estimated that individuals from population cohorts carried a mean of 53 variants previously thought to be sufficient causes of Mendelian diseases. Additionally, 88% of such variants had MAF>1%, implying that they are likely not sufficient causes. This may indicate that such variants are not causally related to disease, or perhaps, that they are causally related but with reduced penetrance.

Penetrance plays an important role in understanding disease pathology, in the appropriate classification of pathogenic variants, and perhaps above all in the context of genetic counseling. However, most of the variants reported to date have been very rare and identified in small sets of unrelated individuals (sometimes just one) or small pedigrees. Penetrance cannot be estimated from a single case, or a single parent-offspring trio presenting with a *de novo* mutation in the offspring. But even with multiple cases or families, determination of the penetrance can present challenges. Here we focus on one such challenge: ascertainment.

Typically a variant of interest is first identified in one individual with a given phenotype. Investigators may then sequence either additional relatives of the individual, or additional individuals or families presenting with the same or closely related phenotypes, with the goal of bolstering the case for pathogenicity. Thus, ascertainment of individuals to be sequenced typically proceeds in stages. The precise ascertainment process used to enrol individuals and/or families is usually at least to some extent unsystematic, and may vary between families. Ascertainment is therefore challenging to model when attempting to estimate the penetrance of a variant.

One situation in which ascertainment can be easily handled is “single” ascertainment, in which the probability of an affected individual being ascertained is proportional to the number of affected individuals in the family (5). In fact, much of the literature on inferring pathogenicity or estimating penetrance tends to assume single ascertainment, e.g., (6), where ascertainment is addressed by conditioning on “the proband,” a procedure which is strictly correct only under true single ascertainment. While it is true that the typical study ascertains families through one individual who may be designated as the single “proband”, this does not ensure that the study meets the proportionality requirement of single ascertainment. This requirement would be violated, e.g., if families with four affected members were more than twice as likely to be recruited as families with just two; or, if the probability of a second sibling being ascertained were dependent on the ascertainment status of the first. And in general, if either (i) ascertainment is not truly single, or (ii) even if it is, if an appropriate ascertainment correction is not incorporated into the estimation method, then penetrance estimates will be biased. Here we consider the magnitude of that bias, across a range of plausible ascertainment models and varying amounts of available data.

We focus here on sibship data. The impact of ascertainment for more complex pedigrees can be approximated by considering large sibship sizes. For simplicity, we assume all parents are phenotypically and genotypically unknown; including parental information does not substantively affect results. We assume a very rare variant of interest (VOI), and an autosomal dominant disease D. Let a qualifying individual (QI) be anyone who is both heterozygous (HET) for the VOI and also affected (AFF) with D. Let *r* be the number of QI sibs within a family, and let *t* be the number of AFF sibs regardless of VOI genotype. We also assume that, regardless of VOI status, an individual might develop D due to other factors, which might be genetic (involving one or more VOIs at other loci or other variants within the same gene) and/or environmental (e.g., due to infections). Let *γ* be the combined penetrance across all causes other than the VOI under study. Since we assume the VOI is very rare, *γ* is effectively the population prevalence of D.

In order to consider a range of plausible ascertainment scenarios, we employ the general family-based *k*-model of ascertainment (7). In its simplest form, this model stipulates that the probability that a family is ascertained is proportional to *r^k^*, where *k* controls the model. For example, when *k* = 1, the probability of ascertainment is strictly proportional to *r*: this is equivalent to classical “single ascertainment”. Similarly, when *k* = 0, so that every family with *r* ≥ 1 is ascertained, this model is equivalent to classical “complete” or “truncate” ascertainment. We generalize this model in two ways. First, we assume that ascertainment requires *r* ≥ 1, that is, every ascertained family contains at least one QI, but we allow that there may be additional preferential ascertainment of families based on *t* alone, that is, that investigators may preferentially ascertain families with more affected individuals without knowing (or prior to knowing) the VOI status of those additional individuals. Second, we allow that even an individual carrying the VOI may develop disease due to any other independent causes at work in the general population. With these two extensions in mind, our ascertainment model becomes

P[sibship is ascertained | *r*, *t*] = *c*(*r^k^* + *t*); *for r* ≥ 1, and 0 otherwise

where *c* is a normalizing constant.

Let *f* be the attributable penetrance, or the penetrance due to the VOI for HET individuals. (Note that when *γ* > 0, *β*=P[AFF|HET]= *γ* + *f* – *γf*. However, we focus here on estimation of *f* itself rather than *β*.) In what follows, we estimate *f* in two ways:

(ii) 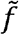 is obtained by counting the proportion of AFF individuals among all HET individuals in the data set, after dropping one QI individual per family, that is, applying the correction for single ascertainment;

(ii) 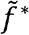 is obtained by counting the proportion of AFF individuals among all HET individuals in the data set, that is, without applying any ascertainment correction.

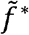 is a naïve estimate, which would be correct if the families were not ascertained based on either phenotype or genotype. It is, however, clearly incorrect under any of our ascertainment models. Our interest in this estimate is to establish how biased it becomes under various ascertainment scenarios. 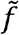 by contrast, does apply the frequently employed single ascertainment correction, and again, our interest in 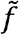 is to establish how biased it will be under ascertainment scenarios other than single ascertainment. Expected values of and 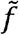 and 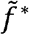 were obtained via simulation, by averaging each estimate’s value across 1,000 replicates per generating condition, and standard errors were obtained by averaging the standard deviation of each estimate across those same 1,000 replicates. (While the expected values are easily calculated analytically, the standard errors are not.) All simulations and calculations were done in MATLAB (2021.9.10.0.1739362 (R2021a), Natick, Massachusetts: The MathWorks Inc.).

Let *s* be the number of siblings in a family, and let N be the number of *s*-sized sibships in a dataset. Fig 1 shows results for true single ascertainment (k=1), for *s* = 2, as a function of sample size N. Here we assume that the true value of *f*=0.5. As can be seen, in this case, the mean of 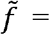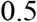, the generating value, as expected. But using 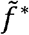 the estimates are seriously upwardly biased in all data sets, regardless of N. Note that because each sibship contains at least one QI, by stipulation, the minimum value of 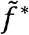 is 0.50.

**Figure 1.**
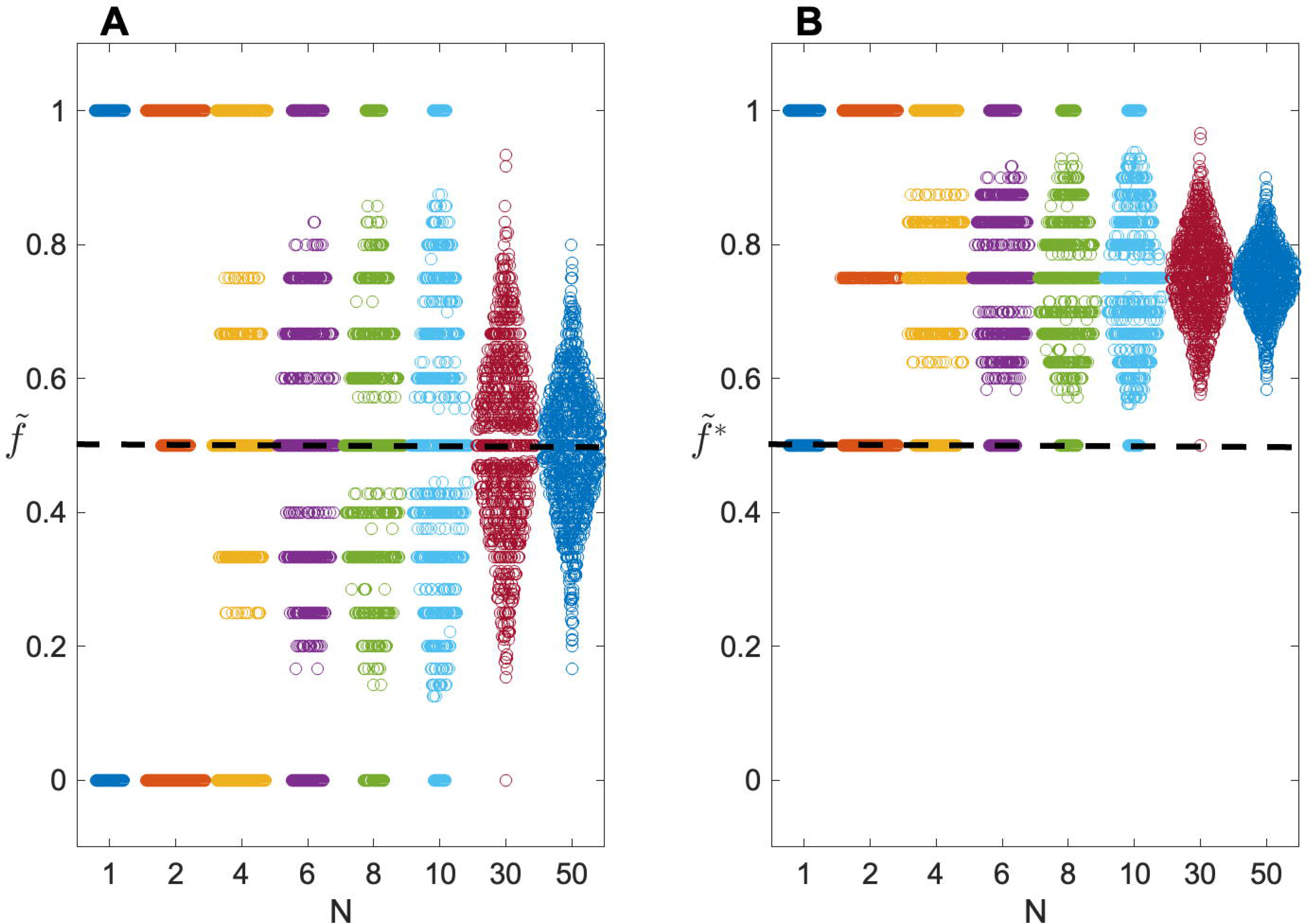
Swarm plots showing sampling distributions of penetrance estimates as a function of number of families N. Distributions of (A) 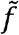 and (B) 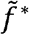 are shown for simulations of 1000 replicates, with true penetrance *f*=0.5. The number of sibs per family, s=2; phenocopy rate, *γ*=0. Users interested in varying the parameters can use the PenEst app.

Note too that even the correct estimate 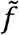 shows considerable sampling variability. For instance, with N=10, 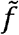 will be >70% or <30% in approximately 40% of all data sets when *f* =50%. This variability remains appreciable even for N=50.

For ascertainment models other than single, overall variability remains similar to what is shown in Fig 1, but even 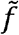 tends to be biased, with mean 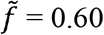, 0.50, 0.43 and 0.38 for *k* = 2, 1, 0 and −1, respectively. In all cases, the uncorrected 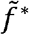 will return even more biased estimates, with mean 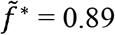, 0.88, 0.87 and 0.86, for *k* = 2, 1, 0 and −1, respectively.

Fig 2 shows the impact of the population prevalence *γ* on average penetrance estimates. Focusing first on single ascertainment (*k*=1) and 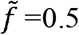, we can see that regardless of *k*, the expected value of 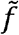 is relatively independent of *γ* until *γ* becomes quite high. Note that for *f* = 0.5 and *γ* = 0.5, the actual probability that a VOI carrier is affected under our generating model is 0.5 + 0.5 − (0.5)(0.5) = 0.75, which is in line with the estimates returned by 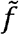. 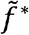 might be said to be even more robust to *γ*, although this is because in this case 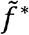 is already close to the top of the scale for *γ*=0. Moreover, 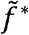 appears not only robust to *γ*, but also to *f* itself, with estimates >70% even for *f*=0.05, and >80% for *f*=0.05 when *γ*=0.5. These patterns repeat for different values of *k*, with visible impact only on the magnitude of the bias for any given (*f*, *γ*) combination. Ascertainment effects will be reduced as *s* increases. Users who are interested in investigating penetrance estimates for other ascertainment models, other combinations of parameter values or other sibship sizes are encouraged to download the PenEst app: https://github.com/MathematicalMedicine/PenetranceEstimator.

**Figure 2.**
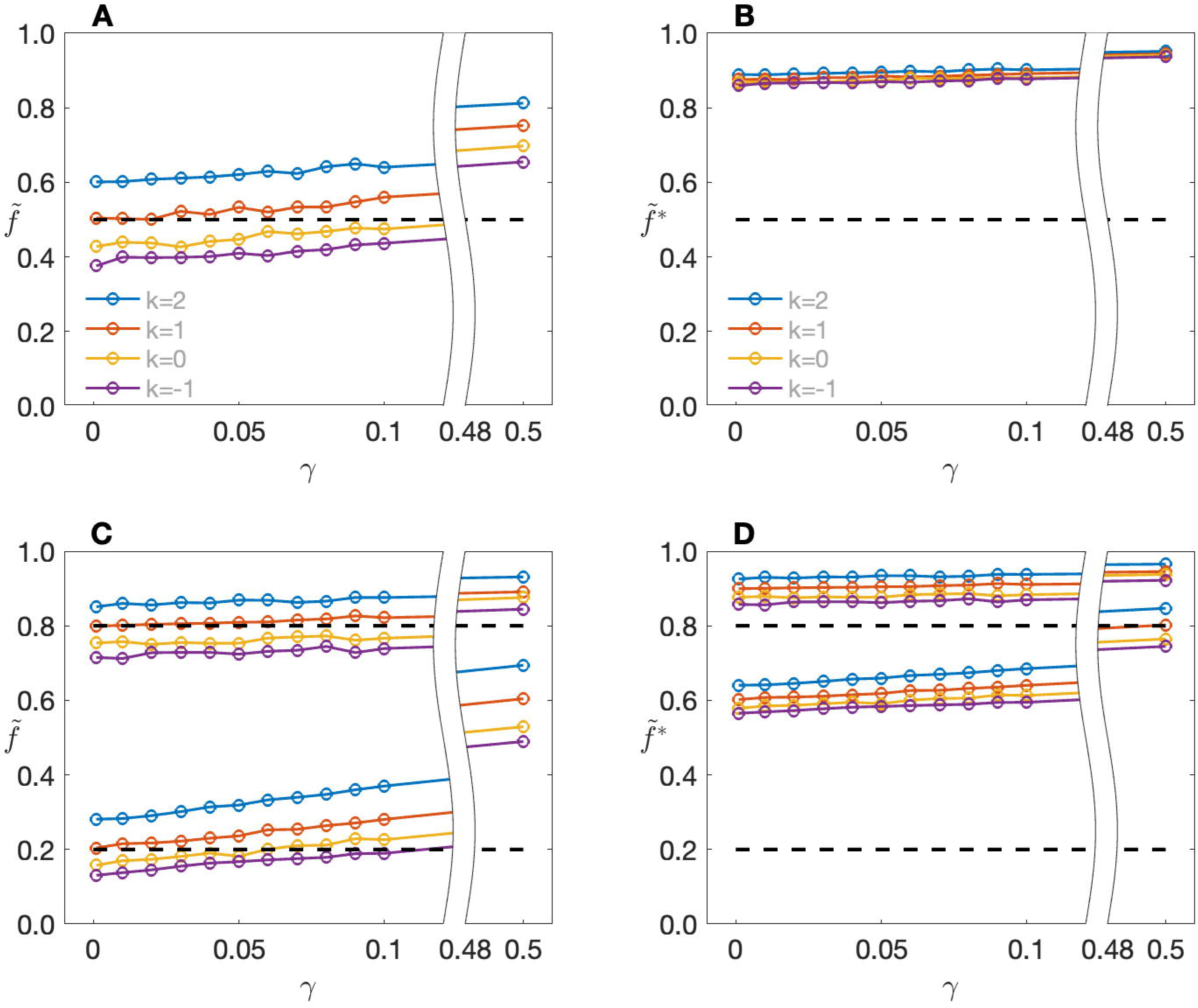
Expected values of penetrance estimates as a function of population prevalence *γ* and ascertainment parameter *k*. Top row: expected values of (A) 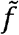 and (B) 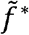 when the true penetrance *f*=0.5. Bottom row: expected values of (C) 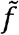 and (D) 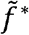 when *f*=0.2 (lower line sets) or *f*=0.8 (upper line sets). The number of sibs per family, s=2. Users interested in varying the parameters can use the PenEst app.

In general, our simulations show that under unsystematic ascertainment schemes, or in cases where appropriate ascertainment corrections are not included in the estimation procedure, there is a high risk of over-estimating the penetrance of any given VOI. This finding is consonant with, and may in large part explain, reports for specific variants. For example, multiple coding variants in *PRNP* had been reported to cause rare dominant monogenic neurodegenerative disease, but there was a 30-fold higher prevalence of variants previously suggested to be causal in this gene in ExAC compared to the expected frequency calculated from the estimated prevalence of the disorder (8). Specifically for three variants the lifetime risk of developing disease was <10%. Similarly, GWAS array data from the UK Biobank were used to estimate pathogenicity, penetrance, and expressivity of putative disease-causing rare variants (MAF<1%) that were directly genotyped and had good quality (9). Focused on maturity-onset diabetes of the young and developmental disorders, many specific variants were found for which the penetrance -- estimated either in families ascertained for the presence of the VOI or in disease cohorts -- was much higher than that obtained from a population-based cohort. These observations have implications for genetic counselling, including the recommendation of invasive screening procedures and administration of preventative treatment.

Some approaches to the interpretation of rare coding variants assume either full or high penetrance (10), for the sake of simplicity. Extensive criteria have been proposed to claim a causal relationship between variants and disease, and authors have urged caution in presuming full penetrance for pathogenic variants (11). But in practice, penetrance remains an important factor in assessing pathogenicity. For instance, the ACMGG/AMP joint consensus recommendations (1) warns against ignoring the possibility of reduced penetrance in establishing segregation of a VOI with a phenotype, but also instructs that “lack of segregation…provides strong evidence against pathogenicity.” (p. 15) And in practice, many laboratories will rule out candidate VOIs when they are found among unaffected relatives. Particularly in the absence of a rigorous and accurate estimate of the actual penetrance, this complicates the use of segregation information in assessments of pathogenicity.

We close by noting that there is one essentially “ascertainment assumption free” (12) method for estimating the penetrance, viz., by conditioning on all of the phenotypic data. This is the ascertainment correction implicit in the usual LOD score (13–15), and also the LOD score allowing for linkage disequilibrium or LD-LOD (6, 16, 17), and in principle any program that allows calculation of the LOD score will support this method. As in Thompson (6) the calculation is done here assigning the VOI (which plays the role of the “marker”) and the disease allele the same (rare) frequency (we have used 0.001 in the simulations), assuming complete linkage disequilibrium between the two (D’ = 1), and also assuming 0 recombination between the marker and the disease allele. Free parameters in the model are then the three penetrances; in our calculations we also include the admixture parameter *α* of Smith (18), representing the probability that any given family is of the “linked” type, which adds robustness when phenocopy levels are high. Maximizing the LD-LOD over the free parameters gives us the LD-MOD, which occurs at the maximum likelihood estimate (m.l.e.) of 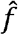 of *f* (12–15).

Fig. 3 shows results corresponding to the simulations in Fig 1A and Fig 2A, C. As can be seen, 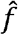 behaves very much like 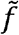 when k = 1 (Fig 3A), but it retains almost complete robustness to ascertainment, and also to *γ* at least until *γ* is quite large (Fig 3B). (As with 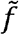, as *γ* gets very large, 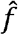 covers both cases due to the VOI and also cases among variant carriers due to other causes.) Comparing Fig 3A with Fig 1A, 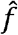 shows slightly greater sampling variability than 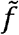; this is due to the inherent ascertainment correction built in to 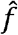. The slight but systematic over-or under-estimation of *f* seen in Fig 3B is due to the small sample size; as N increases 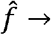 f (results not shown). However, in small samples the upward bias can be appreciable particularly when *f* is small; e.g., when *f* = 0.05 (y = 0), for N = 20, the expected value of 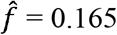.

**Figure 3.**
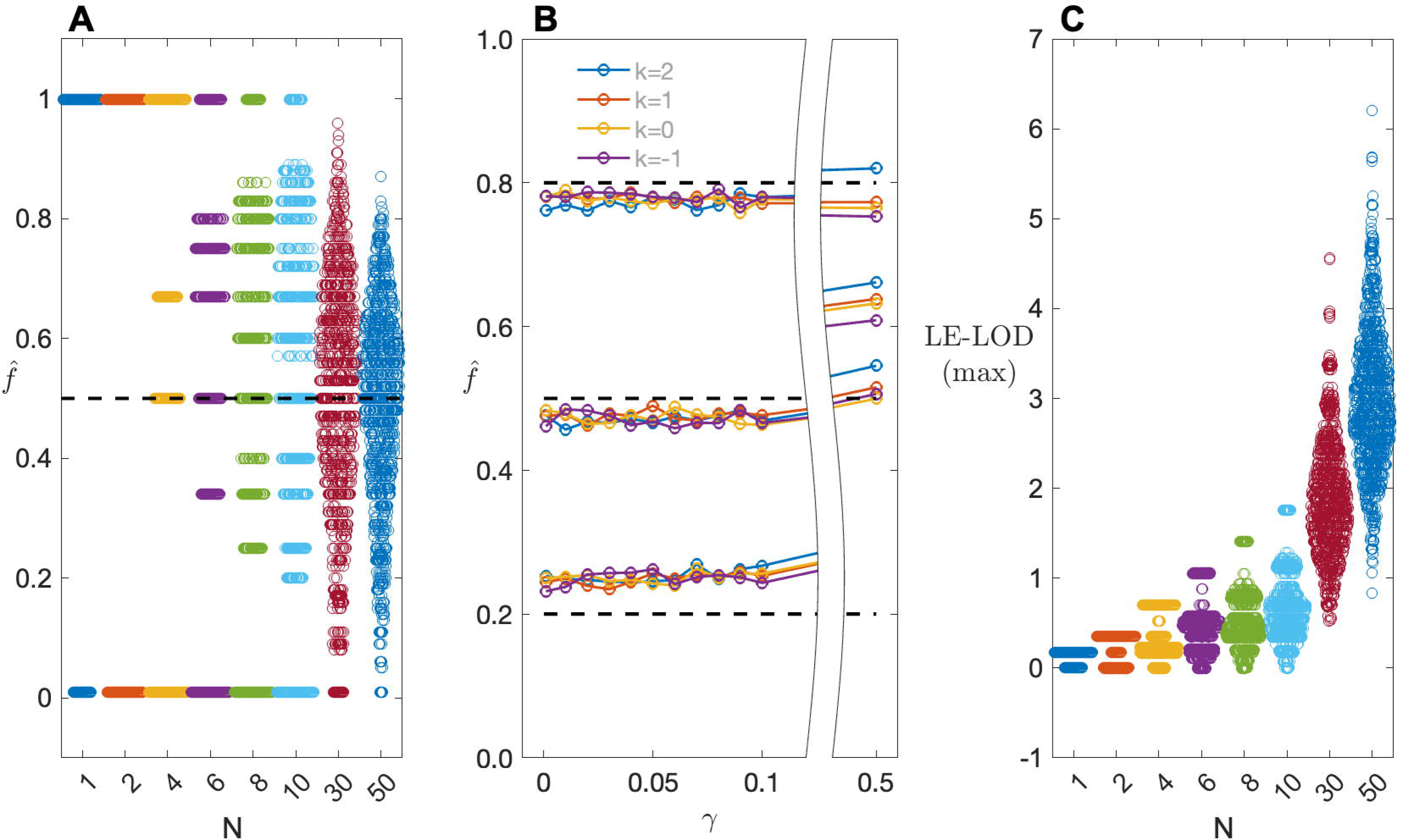
(A) Swarm plots showing sampling distributions of 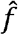, as obtained from maximizing the LD-LOD, as a function of number of families N; (B) Expected values of 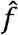 as a function of population prevalence *γ* and ascertainment parameter *k*, for *f* = 0.2, 0.5 and 0.8, reading from bottom to top of the plot, respectively; (C) Swarm plots showing the distribution of LE-LOD(max) as a function of N. Data are the same as used to generate Figures 1 and 2, respectively. All calculations were done using KELVIN (19).

Note, however, that while maximizing the LD-LOD is a highly ascertainment-robust method for estimating *f*, the LD-MOD itself is not a good statistic for representing the strength of evidence for co-segregation, because it is not additionally conditioned on ascertainment through the VOI. However, once we ascertain so as to require the VOI to be present in the family, there is no remaining LD information in the sibship, since LD information is conveyed entirely by the marker allele frequencies in the parents. Therefore, we recommend using the ordinary (linkage equilibrium) LOD, or LE-LOD, for assessing strength of evidence for co-segregation. Because maximizing the LE-LOD itself will not return true m.l.e.s of *f* under the LD model, we recommend evaluating the LE-LOD at the maximizing model obtained from the LD-MOD, for a statistic we annotate as LE-LOD(max). (This maximization procedure is not inherently inflationary; see Supplemental Results (A). Thompson et al. (6) proposed a form of Bayes factor for assessing evidence for co-segregation of the VOI with disease; see the Supplemental Results (B) for some comparisons between their Bayes factor and LE-LOD(max).) Fig 3(D) shows the distribution of the LE-MOD(max), for the same data shown in Fig 1. (Here parents are treated as genotypically known but phenotypically unknown.) As expected, based on just a few 2-child sibships, evidence of co-segregation of the VOI with disease is quite weak. It requires at least N = 30 2-child families before there is a reasonable chance of obtaining a substantial LE-LOD(max).

## Acknowledgements

This work was supported in part by Mathematical Medicine LLC, with special thanks to Jo Valentine-Cooper for creation of the PenEst app.

## Web resources

PenEst app : https://github.com/MathematicalMedicine/PenetranceEstimator/

## Data and code availability

https://github.com/MathematicalMedicine/PenetranceEstimator/

