## Supplementary figures and images for "The effect of ascertainment on penetrance estimates for rare variants: implications for establishing pathogenicity and for genetic counselling"

### Supp Fig 1

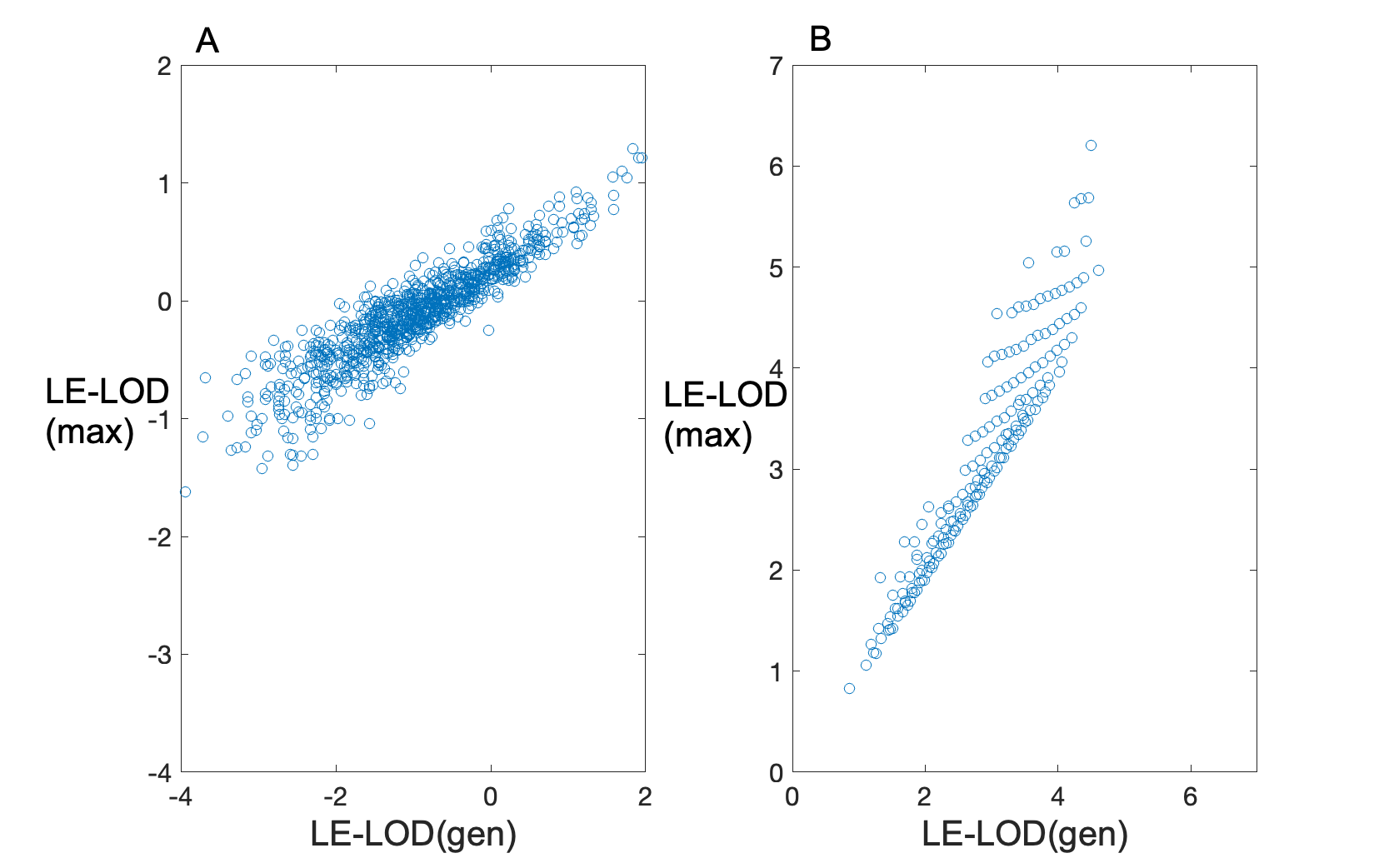

### Supp Fig 2

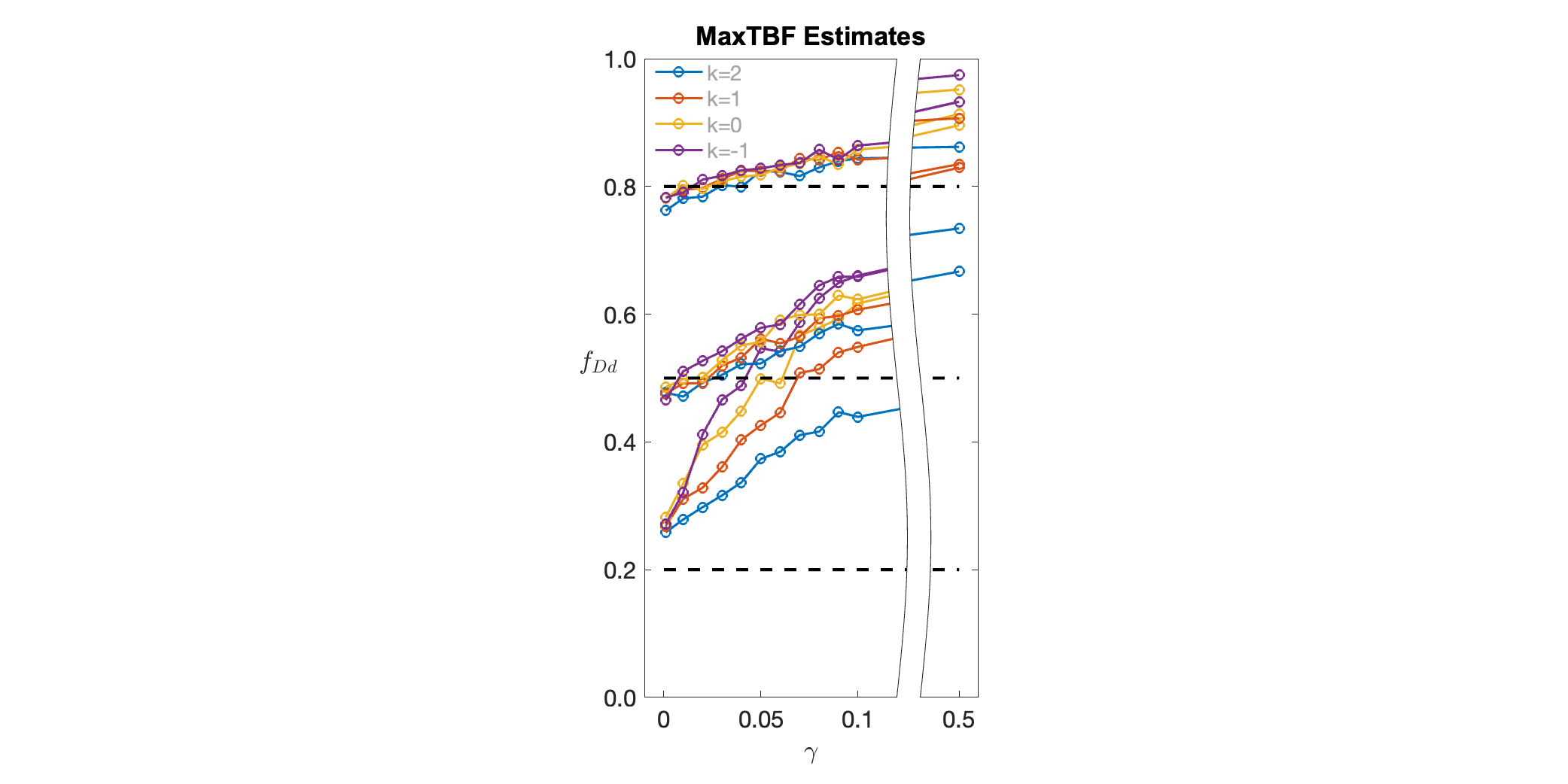

### Supp Fig 3

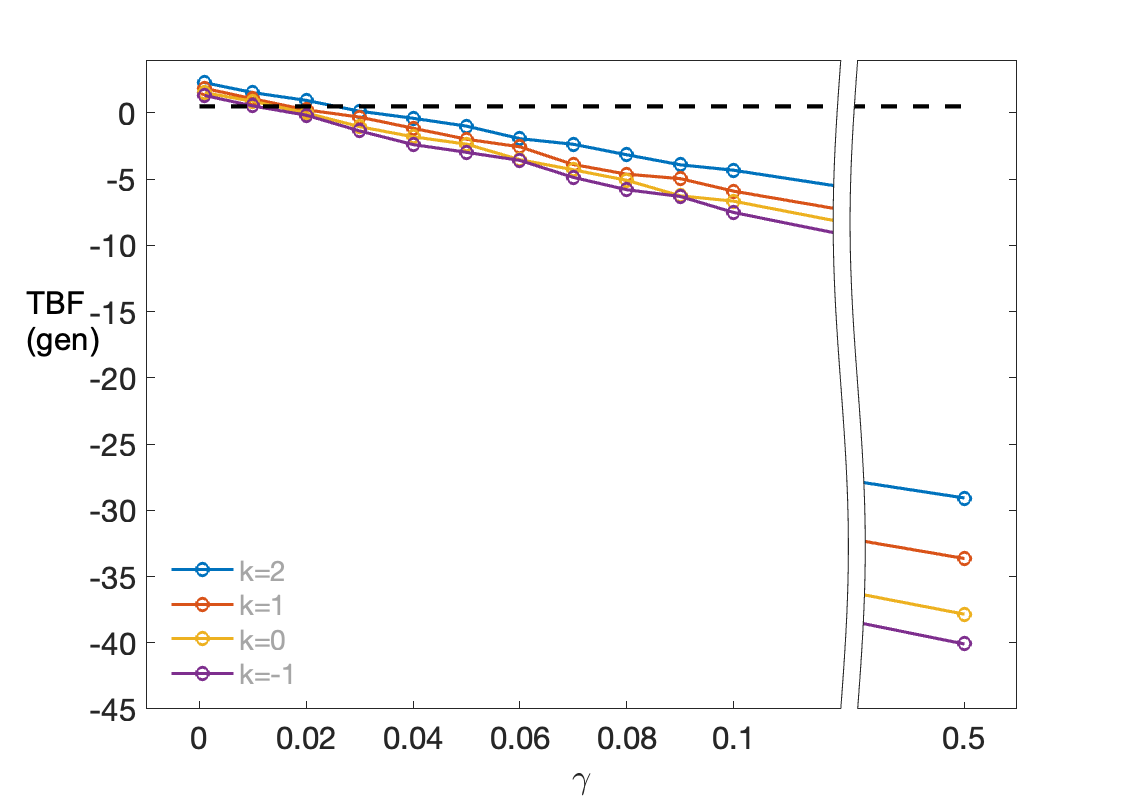

### Supp Fig 4

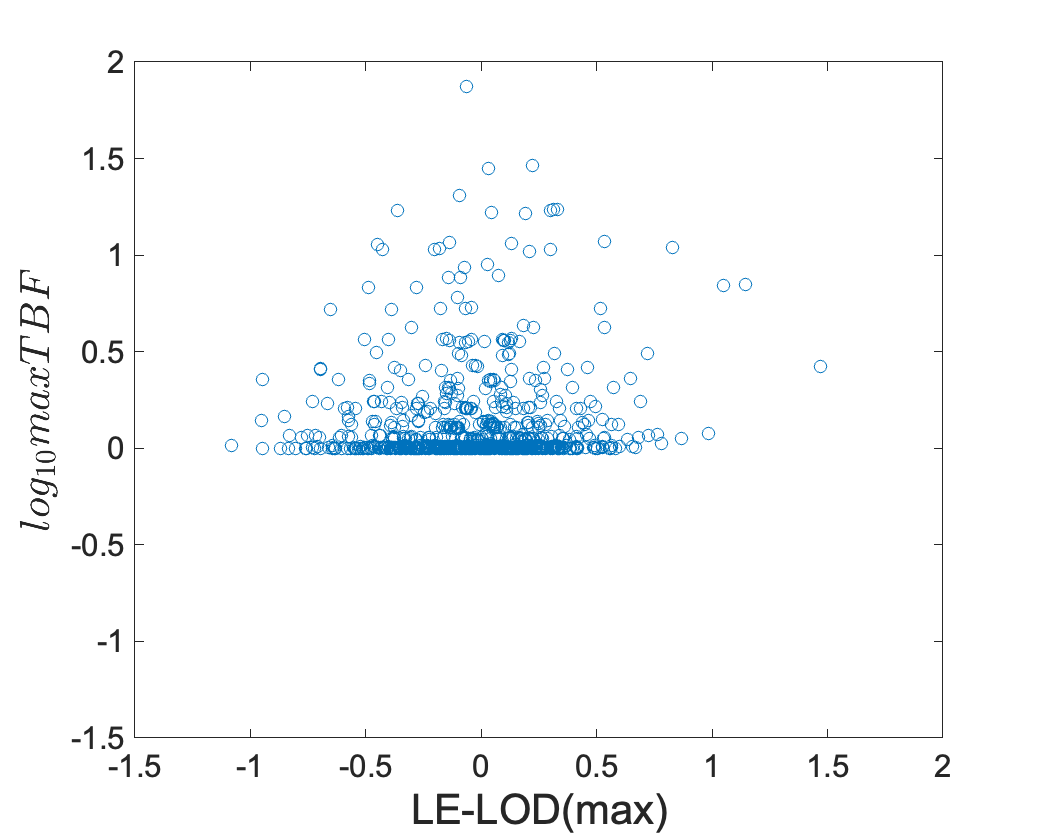

### Supp Fig 5

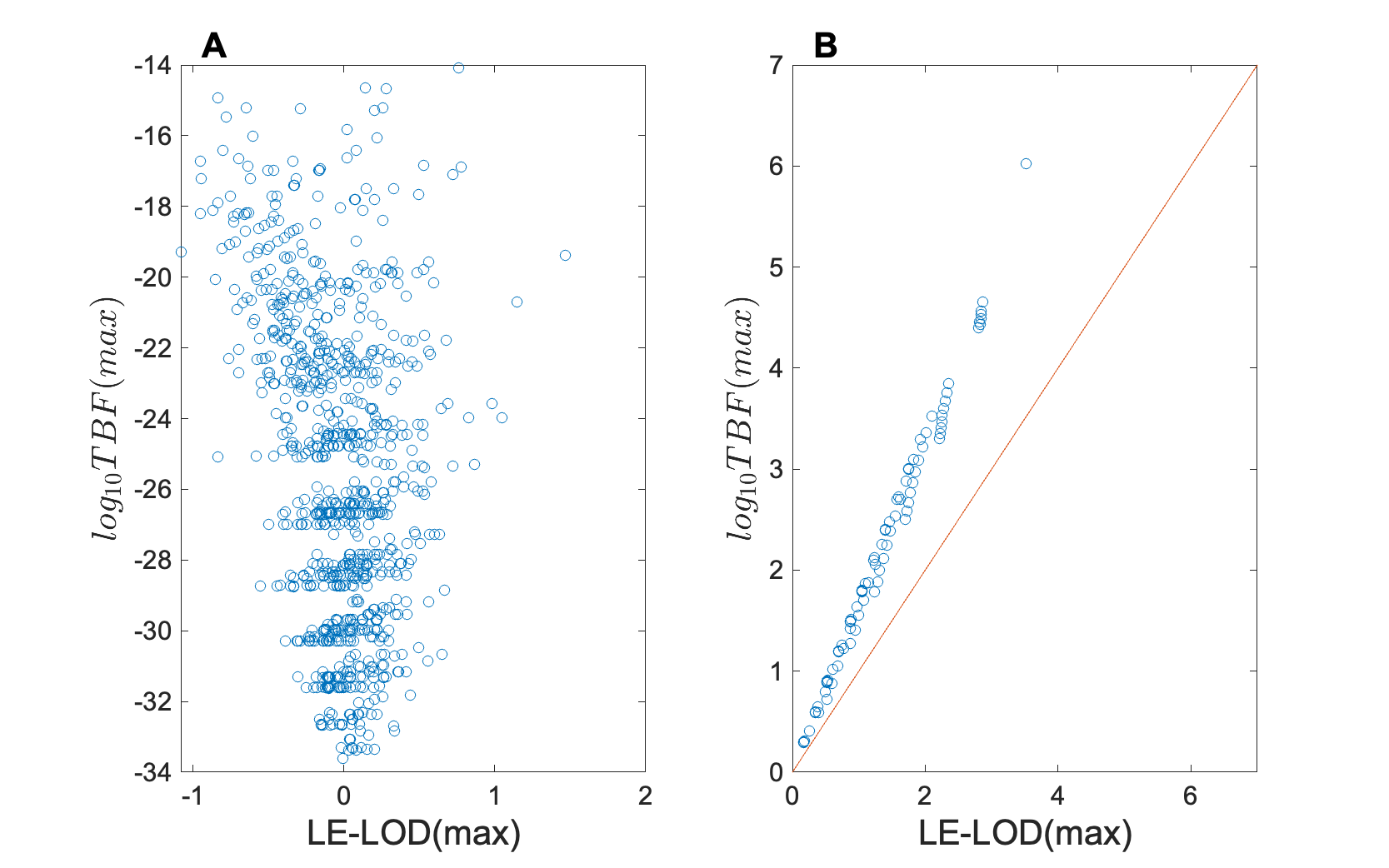
