## Supplementary Materials for "The effect of ascertainment on penetrance estimates for rare variants: implications for establishing pathogenicity and for genetic counselling"

**Supplemental Results**

(A) **Use of the LE-LOD(max) is not inherently inflationary**

Under the hypothesis of no LD and no linkage (no co-segregation of the VOI with disease), the LE-LOD(max) (that is, the LE-LOD evaluated at the trait model estimated by maximizing the LD-LOD) is highly correlated with LE-LOD(gen) (that is, the LE-LOD calculated at the true underlying trait parameter values), as seen in Figure S1A, so that the chance of observing LE-LOD(max) > 3 when LE-LOD(gen) < 3 is negligible. In fact, LE-LOD(max) tends to be smaller, on average, than LE-LOD(gen), with 92.6% of replicates showing LE-LOD(max) < LE-LOD(gen). (Recall that LE-LOD(max) is computed at the model obtained by maximizing the LD-LOD, not the LE-LOD.) Under the hypothesis of co-segregation, LE-LOD(max) does tend to be somewhat inflated relative to LE-LOD(gen), as seen in Figure 1B. However, since here there really is co-segregation, and particularly given the slight deflation in LE-LOD(max) relative to LE-LOD(gen) under “no co-segregation,” the slight inflation in LE-LOD(max) actually facilitates distinguishing the two hypotheses. And again, under the alternative hypothesis, the chance of observing LE-LOD(max) > (say) 3 when LE-LOD(gen) < 3 is very low.

**Figure S1** Scatter plots for LE-LOD(max) under the hypothesis of (A) “no co-segregation” (no LD or linkage) and (B) “co-segregation” (LD and linkage), respectively, for 1,000 replicates of data sets of size N = 50, when s=2, *f*=0.5, $\gamma$=0 and *k*=1.

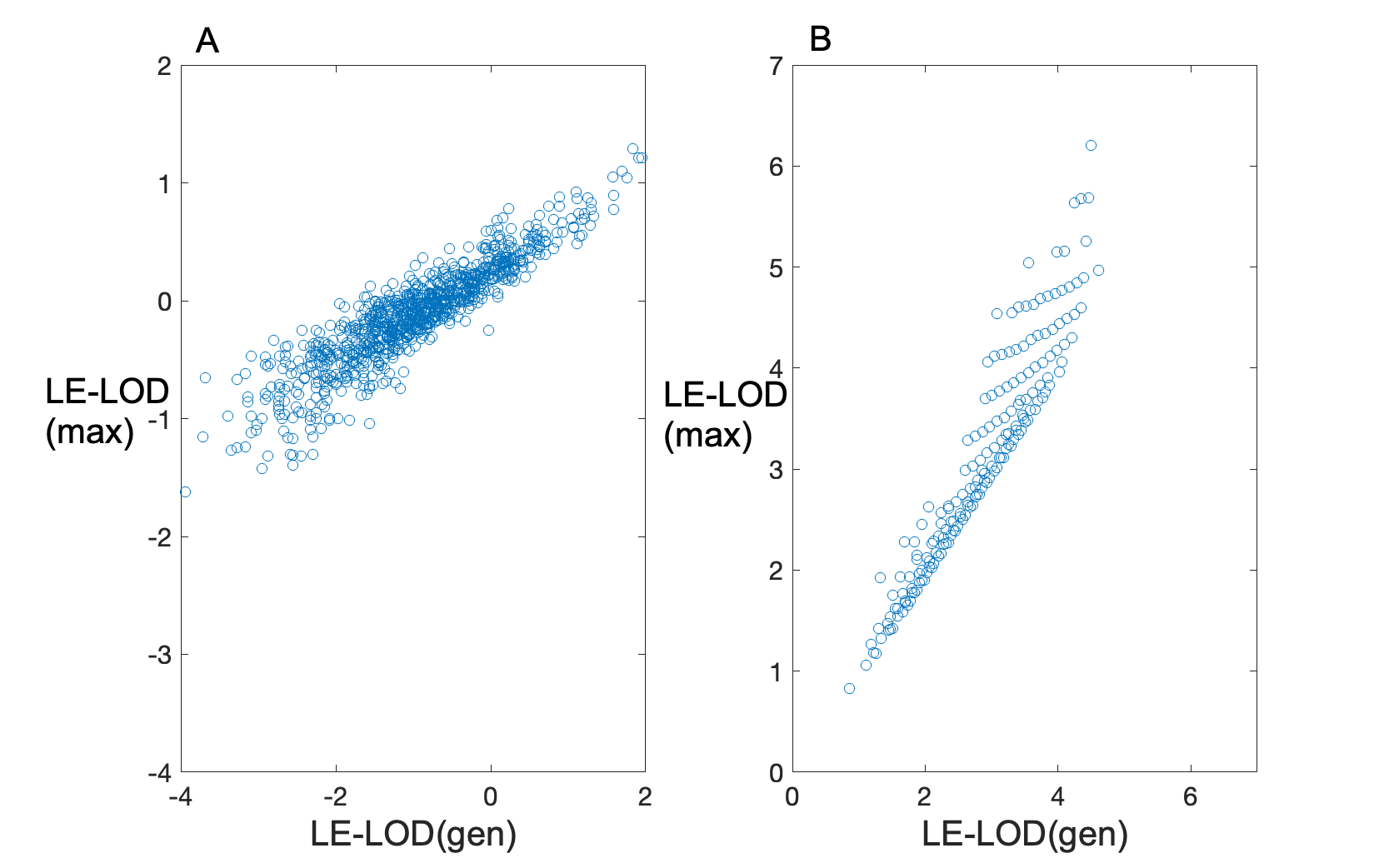

(B) **Comparison with Thompson et al.’s Bayes factor**

Thompson et al. [Thompson D, Easton DF, Goldgar DE. A full-likelihood method for the evaluation of causality of sequence variants from family data. Am J Hum Genet. 2003;73(3):652-5], following [Petersen GM, Parmigiani G, Thomas D. Missense mutations in disease genes: a Bayesian approach to evaluate causality. Am J Hum Genet. 1998;62(6):1516-24], proposed using a particular form of what they refer to as a Bayes Factor (BF) for assessing the strength of evidence for co-segregation of the VOI with disease. This Thompson BF (TBF) is closely related to the LD-LOD discussed in the main text, with an additional adjustment for single ascertainment through a QI (i.e., an individual who is affected and carries the VOI). While Thompson et al. made a compelling case in favor of the TBF, they did not include a systematic evaluation of its operational characteristics, for example, as evaluated using simulations as we do below. Here we hightlight a few salient differences between the behavior of TBF and the LE-LOD(max) we propose in the main text. For comparability with the various forms of LOD scores shown below, we report TBF on the log_10_ scale.

In general, the trait model is not known. One advantage of the LD-LOD is that it is proportional to the likelihood for the marker data conditioned on all of the trait data, an implicitly “Ascertainment Assumption Free” likelihood, as discussed in the main text, and as a result maximizing the LD-LOD over the penetrances is a proper method for obtaining maximum likelihood estimates. The same is not true, however, of TBF. (We note that Thompson et al. did not recommend maximizing the BF to obtain parameter estimates; see also below.) Table S1 below shows results for 2 values of *f* and 2 values of $\gamma$. We have included Smith’s admixture parameter $\alpha$ in the likelihood, as mentioned in the main text, for both LD-LOD(max) and maxTBF. Data were generated with *k* = 1 (the best-case scenario for TBF), with *s* = 2 and N = 1000 (in order to establish large-sample behavior). Shown here are average penetrance estimates across 100 replicates per generating condition, for both the maximum LD-LOD (maxLD-LOD) and for the maximum TBF (maxTBF).

**Table S1. Average penetrance estimates (s.d.) obtained by maximizing either the LD-LOD or the TBF.**

| $\boldsymbol{True \gamma}$ | ***True f_DD_=f_Dd_*** | **maxLD-LOD** | | | **maxTBF** | | |
| --- | --- | --- | --- | --- | --- | --- | --- |
| **0** |  | ***Est’d f_DD_*** | ***Est’d f_Dd_*** | ***Est’d f_dd_*** | ***Est’d f_DD_*** | ***Est’d f_Dd_*** | ***Est’d f_dd_*** |
|  | **0.05** | 0.07 (0.05) | 0.07 (0.05) | 0.0 (0.0) | 0.64 (0.47) | 0.07 (0.05) | 0.0( 0.0) |
|  | **0.5** | 0.50 (0.04) | 0.50 (0.04) | 0.0 (0.0) | 0.74 (0.24) | 0.49 (0.08) | 0.0 (0.0) |
| **0.1** |  |  |  |  |  |  |  |
|  | **0.05** | 0.32 (0.38) | 0.07 (0.05) | 0.0 (0.0) | 0.93 (0.25) | 0.74 (0.26) | 0.72 (0.27) |
|  | **0.5** | 0.77 (0.25) | 0.50 (0.04) | 0.0 (0.0) | 0.93 (0.14) | 0.66 (0.12) | 0.15 (0.03) |

As can be seen, in the presence of phenocopies, estimates obtained from maxTBF are highly non-robust while maxLD-LOD estimates retain their robustness. Indeed, even ignoring the estimates of *f*_DD_ and *f*_dd_, estimates of *f*_Dd_ obtained from maxTBF are also non-robust to ascertainment, as seen in Figure S2, which can be compared with Figure 3B in the main text.

**Figure S2** Estimates of *f*_Dd_ as a function of $\gamma$ and *k*, based on maximizing TBF, for true *f* = 0.2, 0.5 and 0.8, respectively (*s*=2, N=20).

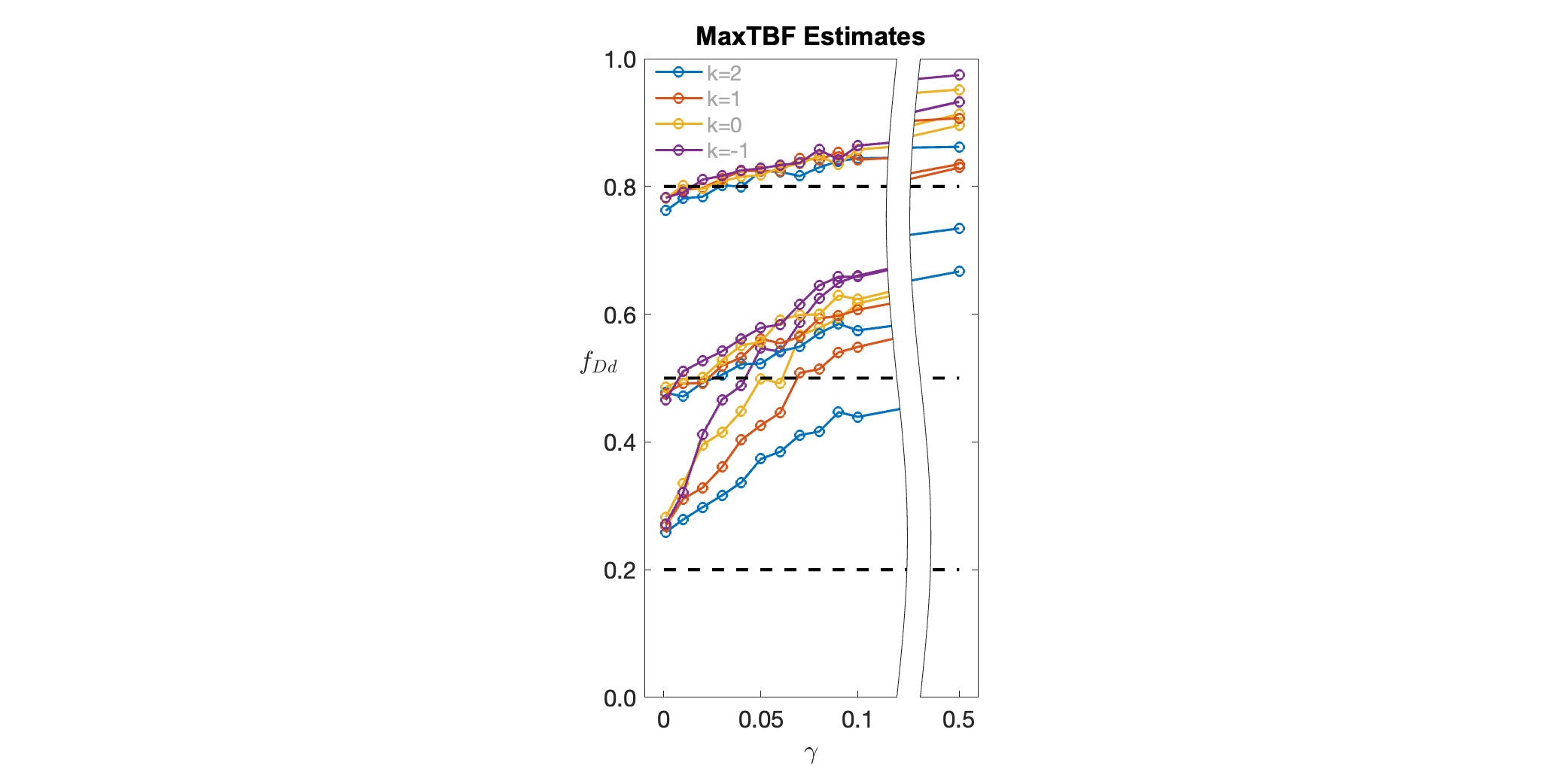

As noted above, Thompson et al. do not suggest using maxTBF to estimate the penetrances; rather, they assume that a reasonable estimate of the penetrances is independently available and can be assumed in computing TBF. However, the behavior of TBF can be quite misleading even if the true model is known exactly. Figure S3 shows the average TBF(gen) (TBF computed at the generating trait parameter values), when the true *f*_Dd_=*f*_Dd_ = 0.5. As can be seen, even with low phenocopy rates, TBF(gen) tends to be negative on average, incorrectly indicating evidence against co-segregation of the VOI with disease.

**Figure S3** Average log_10_TBF(gen) as a function of $\gamma$ and *k* (*s*=2, N=20).

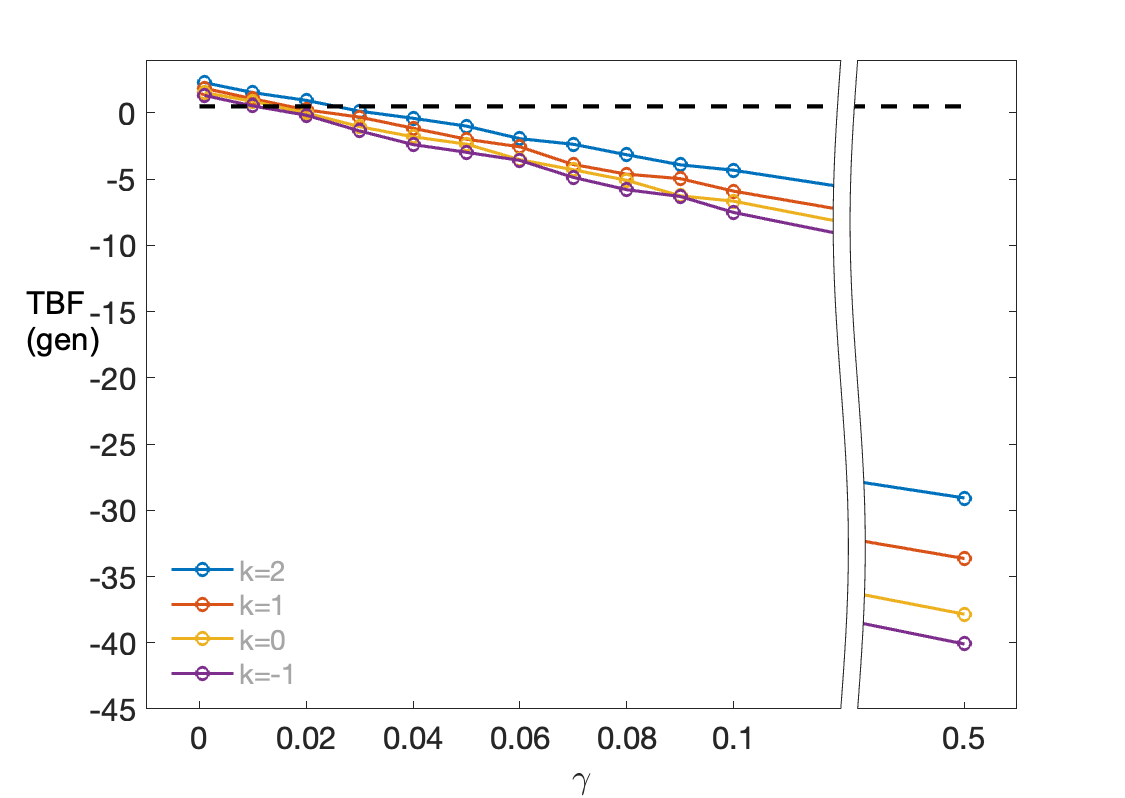

In spite of the tenuous connection between TBF and the true underlying parameter values, it might still be the case that maxTBF performs well for assessing evidence for co-segregation, and the limited results shown here do not settle the matter one way or the other. We note, however, that unlike maxTBF, LE-LOD(max) can in fact indicate evidence against co-segregation, as illustrated in Figure S4. This seems like a critically important feature of any evidence measure.

**Figure S4** Scatter plot of LE-LOD(max) vs. log_10_(maxTBF), for 1,000 replicates (*s*=2, N=20) generated under the hypothesis of “no LD and no linkage.”

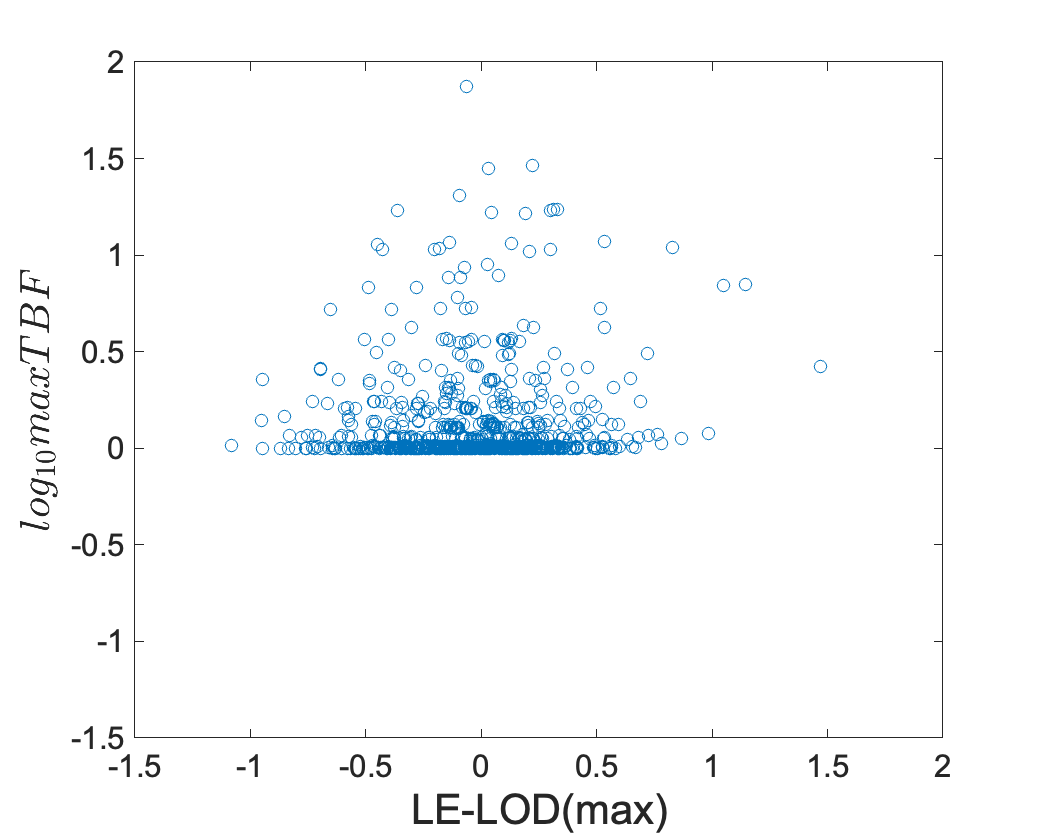

Our final comparison is between LE-LOD(max) and TBF(max), TBF evaluated at the model obtained by maximizing the LD-LOD, that is, the same model used to compute LE-LOD(max). Figure S5 illustrates that the two statistics are quite different in their behavior.

**Figure S5** Scatter plots for LE-LOD(max) and log_10_TBF(max), the TBF evaluated at the same penetrance vector used in computing LE-LOD(max), for data generated under (A) the null hypothesis (“no LD and no linkage”) and (B) the alternative hypotheses (“complete LD and linkage”). Here *s*=2, N=20, *k*=1, *f*_DD_=*f*_Dd_=0.5 and $\gamma$=0. Note in particular the very different scales of the axes in subplot A.

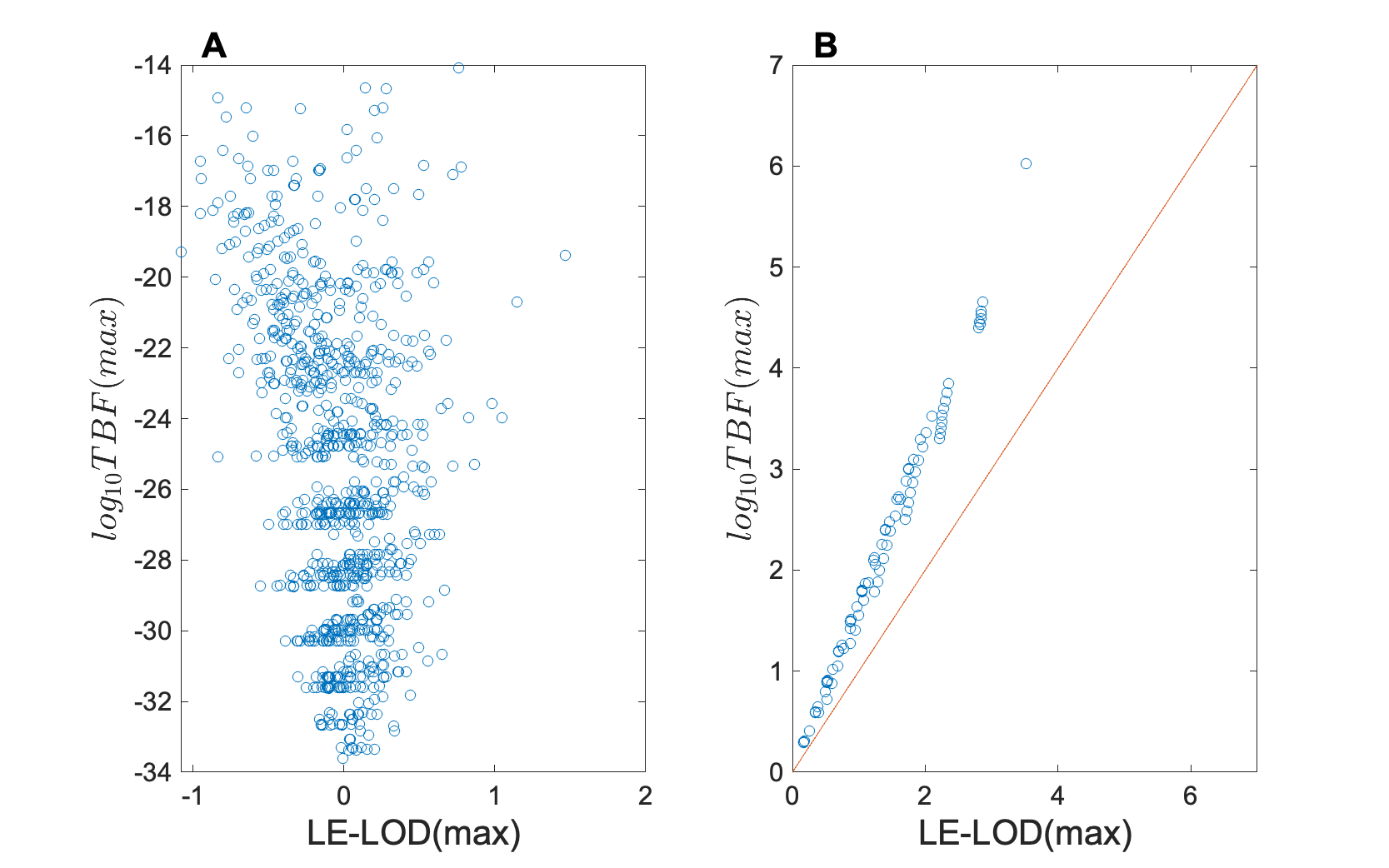

As can be seen, under the alternative hypothesis, log_10_TBF(max) is consistently larger than LE-LOD(max), which might be seen as advantageous. Moreover, under the null hypothesis, log_10_TBF(max) returns strikingly negative results. This might also be seen as advantageous from an operational point of view, in terms of Type 1 and Type II error rates. However, under the null hypothesis, TBF(max) returns numbers that seem wildly out of scale for the quantity of data under consideration.

At the least, TBF(max) appears to be better than maxTBF or TBF(gen). It remains an open question, however, whether having used the LD-LOD to estimate the penetrances, TBF(max) is actually a better or worse indicator of evidence strength compared to LE-LOD(max).
